# Strong spurious transcription likely a cause of DNA insert bias in typical metagenomic clone libraries

**DOI:** 10.1101/013763

**Authors:** Kathy N. Lam, Trevor C. Charles

## Abstract

**Background:** Clone libraries provide researchers with a powerful resource with which to study nucleic acid from diverse sources. Metagenomic clone libraries in particular have aided in studies of microbial biodiversity and function, as well as allowed the mining of novel enzymes for specific functions of interest. These libraries are often constructed by cloning large-inserts (∼30 kb) into a cosmid or fosmid vector. Recently, there have been reports of GC bias in fosmid metagenomic clone libraries, and it was speculated that the bias may be a result of fragmentation and loss of AT-rich sequences during the cloning process. However, evidence in the literature suggests that transcriptional activity or gene product toxicity may play a role in library bias.

**Results:** To explore the possible mechanisms responsible for sequence bias in clone libraries, and in particular whether fragmentation is involved, we constructed a cosmid clone library from a human microbiome sample, and sequenced DNA from three different steps of the library construction process: crude extract DNA, size-selected DNA, and cosmid library DNA. We confirmed a GC bias in the final constructed cosmid library, and we provide strong evidence that the sequence bias is not due to fragmentation and loss of AT-rich sequences but is likely occurring after the DNA is introduced into *E. coli*. To investigate the influence of strong constitutive transcription, we searched the sequence data for consensus promoters and found that *rpoD*/σ^70^ promoter sequences were underrepresented in the cosmid library. Furthermore, when we examined the reference genomes of taxa that were differentially abundant in the cosmid library relative to the original sample, we found that the bias appears to be more closely correlated with the number of *rpoD*/σ^70^ consensus sequences in the genome than with simple GC content.

**Conclusions:** The GC bias of metagenomic clone libraries does not appear to be due to DNA fragmentation. Rather, analysis of promoter consensus sequences provides support for the hypothesis that strong constitutive transcription from sequences recognized as *rpoD*/σ^70^ consensus-like in *E. coli* may lead to plasmid instability or loss of insert DNA. Our results suggest that despite widespread use of *E. coli* to propagate foreign DNA, the effects of *in vivo* transcriptional activity may be under-appreciated. Further work is required to tease apart the effects of transcription from those of gene product toxicity.

## BACKGROUND

Clone libraries can be generated using a range of source material, from the DNA of a single organism to DNA from environmental sources representing often complex microbial communities. Libraries generated from microbial communities are called metagenomic libraries and they have been central to a powerful methodology contributing to understanding the diversity of microbial communities, expanding the knowledge of gene function, and mining for novel sequences encoding functions of interest. These activities all fall under the umbrella of functional metagenomics, and require cloning the DNA, typically using low-copy vectors such as cosmids or fosmids. Cloned DNA is typically propagated in *E. coli* and if the vector host range allows, the DNA can subsequently be transferred to other surrogate hosts that may be more suitable for heterologous expression.

The general assumption in cloning-based metagenomic approaches is that foreign DNA can be stably maintained in *E. coli* and that the cloned DNA is a fair representation of the original sample. However, it has been previously observed that fosmid libraries exhibit a GC bias [1, 2]. In general, such cloning biases may affect conclusions derived from analysis of the clone libraries. The observed GC bias of fosmid libraries was suggested to be due to fragmentation and subsequent loss of AT-rich sequences during the cloning process, purportedly because AT-rich sequences have fewer hydrogen bonds which makes them more vulnerable to non-perpendicular shear forces [1]. Other possible reasons for the bias in libraries include transcriptional activity of the cloned DNA [3] as well as toxicity from expressed genes [4, 5]. Though the exact mechanism(s) by which GC bias occurs has not yet been fully elucidated, the fragmentation explanation has been echoed by others [6, 7] despite being purely speculative and lacking experimental support. Indeed, from our experiences constructing and screening metagenomic cosmid libraries, we believe that events occurring *in vivo* may be contributing substantially to the sequence bias of libraries.

We investigated the nature of this GC bias, to characterize whether, and by what mechanism, biases may be introduced into our own cosmid libraries. In particular, we wished to determine if fragmentation was a major cause of bias, or if there is evidence that the bias was indeed occurring *in vivo*. To answer this question, we constructed a cosmid library using DNA isolated from pooled human fecal samples, saving a portion of the DNA from three steps of the library construction process: (1) the crude extract DNA, (2) the size-selected DNA, and (3) the cloned DNA from the constructed cosmid library (**Figure 1**). The DNA samples were sequenced and the resulting datasets were analyzed to investigate if, where, and how any bias may have been introduced. Consistent with the aforementioned studies, we observed GC bias in our constructed cosmid library. However, our results indicate that fragmentation of DNA does not cause any significant bias; rather, our results are consistent with the hypothesis that the bias occurs after DNA is introduced into the *E. coli* host.

**Figure 1.**
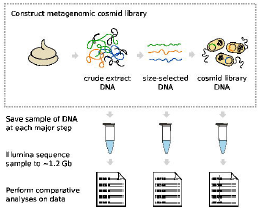
Overview of the experimental design for this study. A pooled human fecal sample was used to construct a metagenomic cosmid library, during which DNA from three distinct steps was collected and sequenced in order to investigate possible sequence biases and at what steps the biases were introduced.

## RESULTS AND DISCUSSION

### DNA sampling and sequencing results

We collected DNA at the three main steps of cosmid library construction: the crude extract DNA, the size-selected DNA, and the final cosmid library DNA (**Figure 1**). Before sequencing, we first checked the quality of each sample by gel electrophoresis (**Figure 2**). As expected, the crude extract was the only sample that contained a heavy smear of fragmented DNA; the selection for high-molecular-weight DNA greatly reduced fragmented DNA, as evidenced by its absence from the size-selected sample. The cosmid library sample exhibited the characteristic multiple banding pattern representing the various possible conformations of uncut circular DNA.

**Figure 2.**
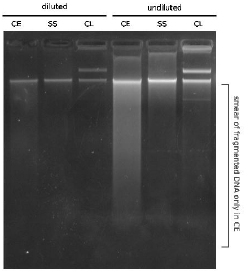
Gel electrophoresis of crude extract, size-selected, and cosmid library DNA samples. Diluted and undiluted amounts of each sample were gel electrophoresed for quality control check of DNA prior to Illumina sequencing.

After confirming DNA quality, the samples were paired-end sequenced on an Illumina HiSeq 2000 platform, generating ∼1.2 Gb of DNA sequence per sample. We expected that the cosmid library would be contaminated with *E. coli* genomic DNA and cosmid vector DNA as a result of (1) isolating cosmid DNA from *E. coli* cells, and (2) the fact that each and every cosmid clone sequenced included its vector backbone. Thus, for fair treatment, we subtracted *E. coli* and pJC8 sequences from all samples (see **Methods**). For *E. coli* and pJC8, respectively: 6 701 and 164 reads were removed from crude extract data (∼0.05% of all reads); 9 273 and 2 410 from size-selected data (∼0.09%); and 851 410 and 2 130 004 from the cosmid library DNA (∼23%). As expected, the dataset originating from the cosmid library sample had the highest number of reads subtracted. Though the crude extract and size-selected samples contained a small amount, these likely represent true environmental sequences; however, their subtraction was necessary for equal treatment of all samples, and the small fraction removed should not affect overall conclusions from the data.

After host and vector sequence subtraction, we used Nonpareil [8] to estimate the overall sequencing coverage of the samples, which was ∼85% for the crude extract and size-selected samples and ∼95% for the cosmid library sample (**Additonal file 3: Figure S3**). This relatively high sequencing coverage was sufficient for our comparative sequence analyses; for all subsequent results discussed in this paper, the forward and reverse sequencing reads for the three samples were analyzed separately.

### GC bias is not caused by fragmentation of AT-rich DNA during cloning

In our experience, extracting high-molecular-weight genomic DNA from low-GC organisms is no more difficult than from *E. coli*. We have previously constructed genomic libraries in cosmid vectors using DNA from *Bacteroides thetaiotaomicron* and *Bacteroides fragilis* (both ∼43% GC) with no difficulties obtaining high-quality DNA [9]. Therefore, it seemed to us that the suggestion by Temperton *et al*. [1] that the GC bias in cosmid/fosmid libraries might be due to fragmentation of AT-rich sequences was unlikely to be true. Our experimental design was such that we could address this specific point: because we sequenced both crude extract and size-selected samples, we could determine whether the removed fragmented DNA from the crude extract (visible in **Figure 2**) led to a bias in the size-selected DNA sample. We examined percent GC in each of the three datasets and found that the GC bias was only present in the final cosmid library and not the size-selected sample (**Table 1**), effectively ruling out fragmentation as the mechanism for cosmid library bias.

**Table 1.** Percent GC of crude extract, size-selected, and cosmid library datasets. GC content was calculated after subtraction of *E. coli* and vector DNA from all samples.

| Sample/Dataset | No. Reads | No. Mb | % GC |
| --- | --- | --- | --- |
| crude extract F | 6 654 484 | 599 | 47.7 |
| crude extract R | 6 654 567 | 599 | 47.8 |
| size-selected F | 6 645 306 | 598 | 46.9 |
| size-selected R | 6 645 817 | 598 | 46.9 |
| cosmid library F | 5 134 020 | 462 | 53.0 |
| cosmid library R | 5 191 538 | 467 | 53.1 |

After confirming that the bias occurs post-size selection, we next asked if certain taxa were differentially represented across the samples to see if this would point to a possible reason for library sequence bias. We used Taxy [10] as well as Taxy-Pro [11] as part of the CoMet web server [12] to do a fast preliminary comparison of taxa abundance across the three different samples. Taxy calculates k-mer frequencies for the dataset and then uses mixture modelling of k-mer frequencies of sequenced genomes to obtain a profile similar to that of the sample, whereas Taxy-Pro has a similar modelling approach but uses protein domains rather than k-mer frequencies. Both tools generated very similar profiles for the crude extract and the size-selected DNA, but a very different profile for the cosmid library DNA (data not shown), supporting the percent GC results. With positive results from this preliminary work, we then performed more thorough taxonomic analyses using two different approaches; in the first, all sequencing reads were used, and in the second, only the 16S rRNA gene-containing reads were used (see **Methods**).

In the first approach, we used MetaPhlAn, a profiling tool that maps reads against clade-specific marker sequences [13], to estimate sample composition down to the species level (**Additonal file 4: Table S4**). We examined the abundance of the top four most common phyla in human gut metagenomes to see whether there were large overall changes in taxa abundance across the samples (**Figure 3**). The crude extract and size-selected samples showed high Firmicutes and

**Figure 3.**
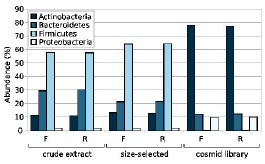
Histogram of abundance of the top four phyla in crude extract, size-selected, and cosmid library samples. Abundance of the Actinobacteria, Bacteroidetes, Firmicutes, and Proteobacteria phyla in each sample, as determined using MetaPhlAn.

Bacteroidetes content with lower levels of Actinobacteria and Proteobacteria, compositions that are typical of gut-derived samples [14–16]. Notably, our results indicated that the cosmid library sample underwent a substantial decrease in the Firmicutes, accompanied by a comparably substantial increase in the Actinobacteria. These results were consistent with the percent GC analysis, as members of the Firmicutes phylum are generally known to be low-GC, and those of the Actinobacteria, high-GC. We also examined the MetaPhlAn results at the species level to see which genomes may be under-or over-represented in the cosmid library, choosing to examine the top 50 most differentially abundant species (**Figure 4**). Several members of the *Bifidobacterium* genus were substantially over-represented in the cosmid library while many members of the Firmicutes were completely or very nearly lost; for example, *Eubacterium rectale, Ruminococcus bromii*, and *Faecalibacterium prausnitzii* were all highly abundant in the original sample.

**Figure 4.**
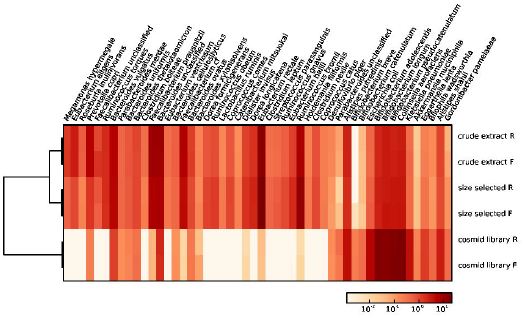
Heatmap of 50 species with differential abundance across crude extract, size-selected, and cosmid library samples. Abundance in each sample of the top 50 species determined to be differentially abundant using MetaPhlAn. Abundance is depicted on a log scale.

In our second approach, we identified reads in the datasets that were from the 16S rRNA gene, and used the RDP classifier to classify these to the genus level (**Additonal file 5: Figure S5**). We observed that analyses using only 16S rRNA gene-containing reads showed high agreement with analyses carried out using all reads (i.e., Figure 4), indicating that 16S rRNA gene content tracks well with genomic content in large-insert cosmid libraries. Both of our approaches provided similar results, and both were in agreement with percent GC, Taxy, and Taxy-Pro results, all of which provide compelling evidence that cosmid library biases are not due to fragmentation of AT-rich sequences during the cloning process.

### GC content may be merely a proxy for *E. coli* constitutive promoter content

From these results, our own previous experiences, and what was previously known in the literature, we had reason to suspect that the cause of the bias occurred *in vivo*. We are not the first to suggest that sequences from AT-rich genomes may resemble the constitutive *E. coli* promoter [17, 18], particularly the –10 Pribnow box. To investigate whether transcription of the insert may be having a negative effect on its maintenance by the host cell, we analyzed the sequence data from the three samples for *E. coli* consensus promoter sequences; in particular, we were interested in examining the data for differential abundance of the *rpoD*/σ^*70*^ consensus sequence, as σ^70^ is the “housekeeping” sigma factor whose promoters are constitutive.

We used the known promoter consensus sequence for *rpoD*/σ^70^ [19], and, as negative controls, we used the consensus sequence for: *rpoE*/σ^24^ [20]; *rpoH*/σ^32^ [21]; *rpoN*/σ^54^, which has a GC-rich consensus [22]; as well as the primary sigma factor of *Bacteroides* species, σ^ABfr^ [23] because the *Bacteroides* genus had comparable abundance across the three samples (**Additonal file 5: Figure S5**) and because *Bacteroides* constitutive promoters are not recognized by *E. coli* [24]. We examined each of the three samples for relative abundance of these five consensus sequences (see **Methods** for details). Our results showed that while the crude extract and size-selected samples had similar promoter content profiles, the cosmid library exhibited a deviation (**Figure 5**). Supporting our hypothesis, only the *rpoD* consensus content was considerably different in abundance, by about an order of magnitude when compared to either the crude extract or size-selected sample. The loss of these specific sequences from the cosmid library suggests that the widely used cloning host *E. coli* may be problematic for cosmid-cloned foreign fragments of DNA that are incidentally transcriptionally active in a constitutive manner, and indeed, these findings are supported by previous reports in the literature, which we discuss in more detail in the following section.

**Figure 5.**
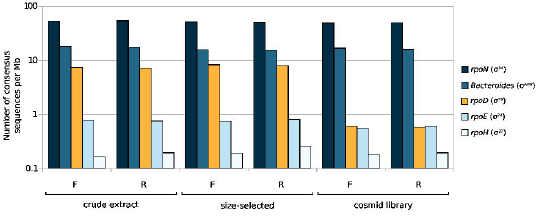
Histogram of sigma factor consensus sequence content in crude extract, size-selected, and cosmid library samples. Bars indicate the number of consensus sequences in each sample, for select *E. coli* sigma factors and the *Bacteroides* primary sigma factor, normalized to the amount of sequence data for that sample. Consensus content is depicted on a log scale.

Given that *rpoD* promoter sequences were under-represented in the cosmid library and that certain species appear to be over-or under-represented, we next asked whether a species' abundance in the cosmid library be predicted from the *rpoD* consensus content of its genome? And in particular, is *rpoD* consensus content more predictive of library abundance than GC content? To answer our questions, we turned to the results of our MetaPhlAn analysis, which gave us a list of the top 50 most differentially abundant species (**Figure 4**). To analyze the genomes of the species for possible sequence determinants of library abundance, we used the NCBI Genome database to find sequenced representatives of each species where possible, and retrieved 46 genomes (complete, draft, or whole genome shotgun sequences; see **Methods** for details); for each genome, we calculated the percent GC as well as the number of *rpoD* consensus promoter sequences present (**Additonal file 6: Table S6**). Next, to quantify bias in the cosmid library relative to the original sample (the crude extract), we calculated the fold change in abundance of the 46 species (using the average abundance of the forward and reverse datasets). We then plotted the fold change in abundance first against genome percent GC (Figure 6A) and second, against *rpoD* consensus content, normalizing to genome size (Figure 6B). Our results show that while library bias only generally correlates with GC content, library bias correlates surprisingly well with the *rpoD* consensus content of the genome.

**Figure 6.**
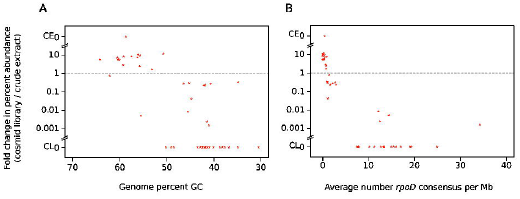
Bias in cosmid library relative to crude extract, against GC content or *rpoD* consensus content. Species abundance was obtained from MetaPhlAn analysis of the crude extract and cosmid library samples. Bias is calculated as fold change in percent abundance (cosmid library abundance / crude extract abundance) plotted against GC content (A) or *rpoD* consensus content (B). Change in abundance is depicted on a log scale; CEo values indicate zero abundance in the crude extract sample and CLo values indicate zero abundance in the cosmid library sample.

These results suggest that GC content may be only a rough proxy for *rpoD* consensus content (as *rpoD* consensus sequences are AT-rich), but GC content itself may not be an accurate predictor of library presence/abundance; indeed, in some cases, a genome may have a moderate or relatively high percent GC but also possess an unusually high *rpoD* consensus content, leading to an underrepresentation in the cosmid library that could not have been predicted from GC content alone (Figure 6). In our view, these results are also consistent with the previous observation that library bias was more obvious among organisms with low GC content [2] because AT-rich genomes would have an increased number of promoter-like sequences simply by chance [25].

### Examining the published literature: evidence for transcriptional activity of cloned AT-rich DNA interfering with stability of circular vectors

In this report, we have presented analysis concerning metagenomic DNA. However, if *rpoD* consensus-like sequences are interfering with the maintenance of foreign DNA in *E. coli*, then the scope of the problem extends beyond metagenomics applications. Curious about the extent of the problem, we performed literature searches to find reports of experienced difficulties cloning AT-rich DNA and/or investigations of possible mechanisms for those difficulties. Our search was fruitful, leading us to literature that spans the past three decades.

It was reported that there are difficulties associated with cosmid-cloning of very AT-rich genomic DNA [26, 27], and even when genomic libraries can be constructed, cosmid clones may be unstable [28–31], which simply means that foreign DNA fragments are not able to be maintained in the *E. coli* library host. Thus, if selection is applied for a marker present on the vector, then *in vivo* events may lead to insert deletion, which has been observed by us as well as others, despite using a host that is a *recA* mutant [31]. This is particularly evident when the library is constructed using a high-copy number vector (e.g., one containing a ColE1-type origin of replication), which has been experienced by us and others [32] and is in agreement with the observation that F-based, singlecopy fosmids perform better than multi-copy cosmids at stably maintaining insert DNA [33]. Loss of cloned sequence is even more widespread for inserts that have repetitive DNA sequences [34], as such sequences may be conducive to recombination. One way to combat insert loss is by minimizing outgrowth of the library-containing cells as much as possible [31], though this is not always feasible for shared cosmid libraries such as our Canadian MetaMicroBiome Library collection [41].

But what is the mechanism for plasmid instability? It was previously shown that transcriptional activity from a cloned strong promoter could affect plasmid stability by (1) interfering with the origin of replication via transcription read-through into the vector, as well as (2) changing the abundance of protein products involved in plasmid copy number. Furthermore, plasmid instability was alleviated by placing transcriptional terminator sequences that flank the multiple cloning site [36]. It was also observed that strong phage promoters could only be cloned into plasmids that possess a downstream termination signal [37, 38]. Similarly, AT-rich pneumococcal DNA was found to contain a high incidence of *E. coli* strong promoter sequences, and that cloning of the DNA was improved by using a vector with efficient transcriptional terminators [3, 32, 39], although analysis of a set of pneumococcal promoter-containing sequences indicated that transcription strong enough to interfere with plasmid stability may be relatively rare and that other factors could be contributing to cloning difficulty [40].

Another consideration is that efficient transcription of poly-dT (as well as poly-dG) DNA tracts may cause the DNA to form a stable complex with its own accumulated transcription products, leading to transcriptional stalling that may interfere with the replication fork [41–43]. One particularly interesting observation that has surprisingly not attracted more interest, is that linear cloning vectors with transcriptional terminators provide even more stability than circular vectors with transcriptional terminators [26, 44]. The advantage of these vectors is due to their linear conformation, but intriguingly, the mechanism remains unclear, although DNA supercoiling of plasmids is thought to play a role (Ronald Godiska, personal communication). These findings along with the aforementioned facts suggest that multiple, distinct mechanisms may be at play to cause cloning bias in *E. coli*, but that there is evidence that transcriptional activity of cloned DNA may be a cause of sequence bias observed in metagenomic libraries. It is often assumed that toxicity of gene products may influence the stable maintenance or “clonability” of DNA in *E. coli* [4, 5, 45] but it is currently unclear whether gene product toxicity is a major factor in the bias of typical clone libraries constructed using circular vectors. It is interesting to consider that cloning bias could be due primarily to purely transcriptional activity rather than the often-blamed protein toxicity.

## CONCLUSIONS

Our own experiences in the lab, the results presented in this report, and what was already known from the literature altogether support the hypothesis that GC bias in typical clone libraries (that is, using circular vectors) is related to promoter activity of the insert in *E. coli*, although DNA topology as well as toxic protein effects may also influence insert and plasmid maintenance. In our analyses, we have focused only on would-be strong constitutive promoters in *E. coli* (sigma 70 consensus sequences) because there is evidence that high level transcription may have negative effects. It is important to acknowledge, however, that functional metagenomic approaches rely on *E. coli* (or other hosts) being able to transcribe and translate foreign DNA, in order to identify fragments encoding functions of interest. This ability of *E. coli* to initiate low-level transcription from diverse sources [46] and to be able to produce foreign proteins, has been immensely advantageous for functional metagenomics, and likely has contributed to the general assumption that *E. coli* is tolerant of foreign DNA, whether it expresses it or not. Our work, however, suggests that more careful consideration of cloning strategies may be required.

Currently, there are three outstanding questions: (1) to what extent does transcription contribute to metagenomic library bias, (2) what factors affect whether transcription will be problematic, and (3) how can transcriptional effects be minimized so that DNA can be faithfully maintained in *E. coli*. An important consideration may be the likelihood of an *rpoD* consensus sequence being cloned on any given fragment from a genome or metagenome. As an example, let us consider *Ruminococcus bromii*, which was one of the most highly abundant species in the original sample but became nearly absent in the cosmid library according to our analyses (∼7% vs. ∼0.01%, respectively; see **Additonal file 4: Table S4**). *R. bromii* has a genome size of 2.25 Mb; theoretically, its genome can be represented in ∼80 fragments if we consider that the average fragment in the particular cosmid library discussed here is ∼28 kb (data not shown). Given that there were 77 *rpoD* consensus sequences identified in its genome (**Additonal file 6: Table S6**), potentially many fragments could include a sequence that behaves as a strong, constitutive promoter in *E. coli*.

In general, it may be helpful to use cloning vectors that include transcriptional terminators flanking the cloning site. We are currently investigating the extent to which transcriptional terminators alleviate the cosmid library sequence bias, which should help tease apart the issue of transcription from that of gene product toxicity. While it is generally recognized that different host backgrounds are needed for functional screening [45, 47–52], it is not as widely acknowledged that the *E. coli* library host itself may be quite limiting. It is interesting that despite decades of using *E. coli* as “the workhorse of molecular biology”, there is still much left to discover about how it tolerates exogenous DNA, which should serve as a reminder to us of how necessary it is to continually re-evaluate even our most basic methodological assumptions, particularly when they concern the inner workings of the cell.

## METHODS

### Sampling of DNA during steps of metagenomic cosmid library construction

Methods for the construction of cosmid libraries, including the specific human gut metagenomic library discussed here (NCBI BioSample ID SAMN02324081), have been previously described in detail [9]. Briefly, DNA was extracted from pooled human fecal samples using freeze-grinding with liquid nitrogen followed by gentle lysis. Crude extracted DNA was then size-selected by pulsed field gel electrophoresis using a CHEF MAPPER Pulsed Field Gel Electrophoresis System (Bio-Rad), followed by electroelution, retaining fragments between approximately 40 to 70 kb. The size-selected DNA was end-repaired, purified, and ligated into the Eco72I site of linearized dephosphorylated pJC8 vector DNA (Genbank accession KC149513). The ligation product was packaged into lambda phage heads using Gigapack III XL Packaging Extract (Stratagene), followed by transduction of *E. coli* HB101. Transductants were recovered on LB agar supplemented with tetracycline (20 μg/ml), and incubated overnight at 37°C. Resulting colonies were enumerated to estimate library size (∼42,000 clones), and colonies were resuspended, pooled, and frozen at -80°C to form the cosmid library stock.

During construction of the cosmid library, DNA was sampled from three steps: (1) the crude extract DNA, (2) the size-selected DNA, and (3) the final cosmid library DNA, prepared from the frozen stock using a GeneJET Plasmid Miniprep Kit (Thermo Scientific).

### Purification, quantification, and Illumina sequencing of DNA

Two of the three DNA samples, the cosmid library DNA and the size-selected DNA, were sufficiently pure for Illumina sequencing, as gauged by 260/280 nm and 260/230 nm ratios (Nanodrop ND-1000 Spectrophotometer); however, the crude extract DNA required further purification. Crude extract DNA concentration was estimated by gel electrophoresis, using bacteriophage lambda DNA as a standard; ∼150 μg in 1 ml was purified and concentrated on the synchronous coefficient of drag alteration (SCODA) instrument (Boreal Genomics), using an established protocol [53].

All samples were re-quantified by gel electrophoresis, using bacteriophage lambda DNA as a standard, and >2 μg of each sample was sent to the Beijing Genomics Institute (BGI, Hong Kong) for 90-base paired-end sequencing on the Illumina HiSeq 2000 platform, using their in-house protocols and reagents for 350 bp fragment library construction. ∼6.7 million reads were obtained in both the forward and the reverse direction, generating ∼1.2 Gb of sequence data per sample. All sequence data have been made publicly available (see **Data** section).

### Subtraction of *E. coli* genome and cosmid vector contamination

The cosmid library sequence data were expected to have substantial contamination with *E. coli* genomic DNA and pJC8 vector sequences. Sequence data were cleaned of contaminating *E. coli* genomic DNA and vector DNA, using BLAT [54] with a conservative criterion of 100% identity. To remove *E. coli* contamination, we used the genome of *E. coli* K12 MG1655 (Genbank accession U00096.3), which to our knowledge is currently the closest sequenced relative of HB101, the library host strain; to remove vector contamination, we used the sequence of pJC8 (Genbank accession KC149513), formatted to simulate Eco72I-cut, cloning-ready vector by removing the 0.8-kb gentamicin resistance gene stuffer present between the two Eco72I sites.

### Taxonomic analysis

To examine taxonomy based on only the 16S rRNA gene sequences present in the data, we identified 16S-containing reads using Infernal version 1.1 [55] and classified them using the RDP Classifier version 2.8 [56]. The classifier output was visualized using the MEtaGenome ANalyzer (MEGAN) version 5.6 [57]. To examine taxonomy using all sequence reads (i.e., not only those identified as 16S reads), we used the Metagenome Phylogenetic Analysis tool (MetaPhlAn) version 2.0, along with its built-in scripts for visualization [13].

### Promoter analysis

To estimate promoter content in the data, we searched for known sigma factor consensus sequences for the *E. coli* sigma factors, *rpoD*/σ^70^ (TTGACAN_15-19_TATAAT), *rpoE*/σ^24^ (GGAACTTN_15-19_TCAAA), *rpoH*/σ^32^ (TTG[A/T][A/T][A/T]N_13-14_CCCCAT[A/T]T), *rpoN*/σ^54^ (TGGCAN_7_TGC), as well as for the *Bacteroides* primary sigma factor, σ^ABfr^ (TTTGN_19-21_TAN_2_TTTG). To do this, we used regular expression pattern matching with Python version 2.7.3; consensus promoter sequences, literature references, and regular expressions are provided (**Additonal file 1: Table S1**).

### Analysis of reference genomes

Genome sequences were downloaded from the NCBI Genbank database as complete genomes, draft genomes, or from whole genome shotgun sequencing projects. Organism names and accession numbers, as well as other relevant information, are provided (**Additonal file 2: Table S2**).

### Data

Raw Illumina sequence data are available at the NCBI Sequence Read Archive under Study SRP031898. Accession numbers for SRA Experiments are: [NCBI:SRX683591] for the crude extract, [NCBI:SRX683589] for the size-selected, and [NCBI:SRX683586] for the cosmid library. In addition, raw data and other relevant data for this study may be accessed online through our website [58].

## LIST OF ABBREVIATIONS

CE,: crude extract;
SS,: size-selected;
CL,: cosmid library;
F,: forward reads;
R,: reverse reads.

## AUTHORS' CONTRIBUTIONS

TCC and KNL designed the experiments. KNL performed the experiments, structured and performed the data analysis, and wrote the paper. TCC provided constructive criticism, revised the manuscript, and provided reagents and materials.

## ACKNOWLEDGMENTS

We thank the seven anonymous individuals who donated fecal samples for the construction of the CLGM1 cosmid library. We are grateful to the Shared Hierarchical Academic Research Computing Network (SHARCNET: www.sharcnet.ca) and Compute/Calcul Canada for computing resources. Research funding was provided by a Strategic Projects Grant (381646-09) from the Natural Sciences and Engineering Research Council of Canada, by Genome Canada for the project “Microbial Genomics for Biofuels and Co-Products from Biorefining Processes” (MGCB^2^), and by a University of Waterloo CIHR Research Incentive Fund. KNL was supported by a CGS-D scholarship from the Canadian Institutes of Health Research.

## ADDITIONAL FILES

**Additional file 1: Table S1.** Consensus sequences for the five sigma factors used, PMID number for the literature reference, and corresponding regular expressions used to search sequence data. (.txt)

**Additional file 2: Table S2.** NCBI Genbank accession numbers for genome sequences of the 46 species selected for GC content and *rpoD* consensus content analysis. (.txt)

**Additional file 3: Figure S3.** Estimate of sample sequencing coverage using Nonpareil. (.pdf)

**Additional file 4: Table S4.** Taxa abundance output from MetaPhlAn for both forward and reverse datasets of each sample. (.txt)

**Additional file 5: Figure S5.** 16S rRNA analysis results using Infernal for identification of 16S-containing reads, RDP classifier to classify reads, and MEGAN for visualization of results. (.pdf)

**Additional file 6: Table S6.** Length, GC content, and *rpoD* consensus content of the 46 genomes selected for analysis. (.txt)

